# The exocyst is an insulin-sensitive regulator of amyloid precursor protein trafficking and amyloid-beta generation in neurons

**DOI:** 10.64898/2026.04.14.717551

**Authors:** Chantell Balaan, Geetika Patwardhan, Rachel Sachs, Hanna Kumasaka, Swasthita Sadagopan, Shion Aou, Amanda J. Lee, Luke T. Nelson, Brian E. Hew, Jesse B. Owens, Noemi Polgar, Michael A. Ortega, Robert A. Nichols, Benjamin Fogelgren

## Abstract

Intracellular trafficking of amyloid precursor protein (APP) critically influences amyloidogenic processing, yet the mechanisms regulating this pathway remain incompletely defined. The exocyst is a highly conserved, insulin-responsive, eight-protein Rab effector complex that directs intracellular transport vesicle targeting and docking. We identified APP in a proteomics screen of neuronal cell surface proteins altered following chemical inhibition of exocyst activity. In SH-SY5Y cells expressing an APP mutant that enhances amyloidogenic processing, RNAi-mediated silencing of exocyst subunits significantly decreased sAPP and Aβ secretion, accompanied by intracellular APP accumulation. High-resolution microscopy revealed APP colocalization with exocyst subunits in the soma and neurites of differentiated SH-SY5Y neurons and primary mouse hippocampal neurons. Live-cell TIRF microscopy demonstrated coordinated movement of fluorescently tagged exocyst and APP proteins. Proximity ligation assays (PLAs) confirmed close (<40nm) APP-EXOC5 association in differentiated SH-SY5Y neurons, primary mouse hippocampal neurons, and histological sections of mouse brain. To examine whether insulin regulates neuronal exocyst activity, as in adipocytes and muscle, we generated SH-SY5Y cells expressing pHluorin-tagged GLUT4. Using this model, we found that chemical inhibition of exocyst activity prevented insulin-stimulated GLUT4 exocytosis to the plasma membrane. Furthermore, PLAs in both primary hippocampal and SH-SY5Y neurons revealed that insulin increased GLUT4-EXOC5 associations while markedly reducing APP-EXOC5 associations. These findings suggest that insulin signaling redirects the exocyst away from APP-positive vesicles towards GLUT-positive vesicles. Together, our data identify the exocyst as a novel insulin-regulated mediator of neuronal APP trafficking and Aβ secretion.

**SIGNIFICANCE STATEMENT:** Alzheimer’s disease is characterized by the accumulation of amyloid-β peptides, which are generated through the processing of amyloid precursor protein (APP). The intracellular trafficking of APP is a critical determinant of amyloid-β production, yet the molecular machinery regulating this process remains poorly understood. Here, we identify the exocyst, a highly conserved protein complex involved in membrane trafficking, as a regulator of APP trafficking and processing in neurons. Furthermore, we demonstrate that insulin signaling influences exocyst activity and its association with APP, revealing a previously unrecognized connection between insulin signaling and amyloid-β generation. These findings provide new mechanistic insights into APP biology and suggest potential pathways linking impaired brain insulin signaling to Alzheimer’s disease.

## INTRODUCTION

A pathological hallmark of Alzheimer’s Disease (AD), the leading cause of dementia, is accumulation of amyloid beta (Aβ) plaques in the brain ^1–3^. The small Aβ peptides that can aggregate over time in the extracellular space arise from amyloidogenic processing of the transmembrane amyloid precursor protein (APP) in neurons. Full length APP is synthesized and transported to the somatodendritic plasma membrane, where the non-amyloidogenic processing pathway begins with cleavage by α-secretase (ADAM10). Alternatively, full-length APP can be endocytosed to the early endosome, where cleavage by β-secretase BACE1 initiates amyloidogenic APP processing ^2^. Following either α- or β-secretase cleavage, the APP C-terminal fragment (either APP-αCTF, APP-βCTF) undergoes translocation from the somatodendritic endosomal system to the axon ^4,5,6^, where it encounters presenilin-containing ɣ - secretase in pre-synaptic termini for a second proteolytic cleavage ^7^. For APP-βCTF, this results in the 40 or 42 amino acid Aβ (Aβ_40_ and Aβ_42_), which is released into the extracellular space, functioning as a positive neuromodulator ^8–9,13^. It has long been observed that Aβ levels in the CNS steadily rise during the prodromal period of AD ^3^, with increasing neurotoxic and synaptotoxic activity, leading to cognitive impairment ^9^.

Understanding the complicated pathways regulating APP intracellular trafficking in neurons is critical to understanding the balance of amyloidogenic versus non-amyloidogenic processing and the risk of developing AD, and may reveal new therapeutic targets for AD. Despite many previous studies in this field, and identification of molecules that do play key roles in the APP trafficking process ^10^, there remain open questions and sometimes conflicting data. Also, findings with new technologies have challenged old paradigms, like the discovery that active γ-secretase is primarily at the presynaptic termini ^11^, and not in the somatodendritic endosomal system, which would then require that APP-CTFs undergo axonal sorting and transport (translocation) for the final proteolytic processing step. Super-resolution microscopy studies have given us a new picture of the processing and sorting of APP, APP-CTF, and sAPP fragments in the endosomal system ^12^, and that APP-CTFs can indeed be selectively shuttled to the axon through the axon initial segment (AIS). Further elucidation of APP trafficking mechanisms in neurons, and identification of signaling pathways that modulate the rate of Aβ synthesis, remain a high research priority for AD.

The exocyst trafficking complex is a Rab effector composed of eight proteins (EXOC1-8), which are conserved from yeast to humans. EXOC6 binds directly to specific Rab GTPases on transport vesicles, and EXOC5 links EXOC6 to the other exocyst subunits as they assemble on the vesicle to aide in transport and docking. Some subunits, like EXOC1 and EXOC7, are known to bind phosphoinositides at the site of exocytosis on the target membrane. The first mammalian exocyst complex characterized was purified from adult rat brains ^14^, but despite decades of exocyst research, we do not have a good understanding of its role in mammalian neurons. It is well established that the exocyst is necessary for the growth cone dynamics and dendrite arborization during neuron development ^15–19^. However, the role of the exocyst in the mature neuron is still unclear. While exocyst subunits have been found at synapses ^20–21^, surprisingly the exocyst does not play a role in neurotransmitter vesicle tethering at the mature synapse ^22^ and does not seem to be critical for synaptic communication ^23^. Reports have shown that several small GTPases associated with exocyst regulation (*e.g.* Rab8, Rab11, Arf6, RalA) also have a role in APP intracellular trafficking and are linked to Aβ generation in AD ^24–27^. However, there have been no previous investigations into a potential role for the exocyst in APP trafficking or Aβ synthesis.

Another intriguing potential exocyst neuronal function could be in the insulin signaling pathways and dynamic regulation of glucose uptake. Twenty years ago, the exocyst was shown to be a necessary mediator for insulin-induced GLUT4 exocytosis in adipocytes ^28^ and we have recently shown this mechanism is conserved in skeletal muscle cells ^29–30^. We hypothesized that this insulin-exocyst-GLUT4 axis may also be important in neuronal metabolism, and if so, the exocyst could act as a molecular mechanism connecting insulin signaling and APP trafficking. In this current study, we report that the exocyst is indeed a critical regulator of both APP trafficking and amyloidogenic processing in neurons, and that insulin signaling modulates this relationship, potentially directing the exocyst away from APP+ transport vesicles and towards GLUT4+ vesicles for GLUT4 exocytosis.

## RESULTS

### Proteomic screen of neuronal cell surface proteins after treatment with exocyst-inhibitor endosidin-2 (ES2) reveals a significant decrease in APP

To quantitatively identify plasma membrane proteins whose exocytosis in neurons is acutely regulated by the exocyst complex, we developed a proteomics assay employing endosidin-2 (ES2), a small molecule that binds to EXOC7 and has been shown to inhibit holocomplex trafficking activity in many cell types ^30,31–33^. For this experiment, we utilized the well-characterized SH-SY5Y human neuroblastoma cells line, which can be differentiated into neuronal cells with an 18-day protocol ^34^. SH-SY5Y were fully differentiated and then treated with 100 µM ES2 or vehicle for two hours, followed by room-temperature biotinylation of exposed cell surface proteins for 10 minutes (**Figure 1A**). Cells were lysed and biotinylated proteins were purified with streptavidin-resin and eluted with elution buffer. Data-Independent Acquisition (DIA) was completed through the IDeA National Resource for Quantitative Proteomics using the Orbitrap Exploris 480 mass spectrometer with 60-minute gradient per sample. Raw DIA data files were analyzed using Spectronaut database search and bioinformatics was completed for quality control, normalization, and differential expression analysis.

**Figure 1:**
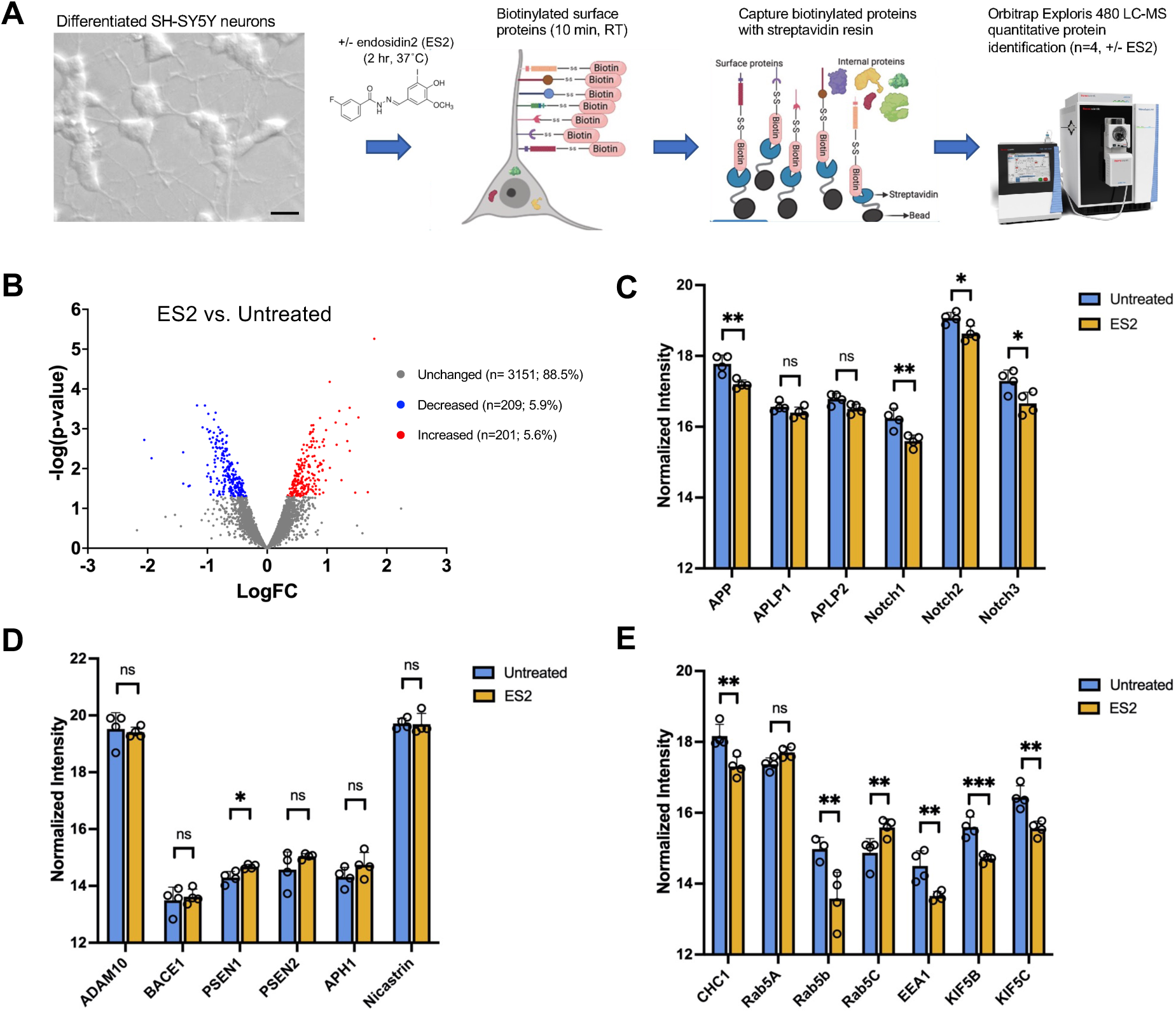
Proteomics reveals reduced amyloid precursor protein (APP) on the plasma membrane of SH-SY5Y neurons when the exocyst is chemically inhibited. **A**. Schematic representation of our experimental workflow. Fully differentiated SH-SY5Y neurons were treated with or without 100 µM endosidin2 (ES2) for 2 hours, surface proteins were covalently labeled with biotin for 10 minutes at room temperature, and biotinylated proteins were then purified with streptavidin columns. Samples (n=4 per treatment group) were acetone precipitated, frozen, and subjected to quantitative protein identification via mass spectrophotometry. B. Volcano plot showing identified biotinylated proteins (3,561 total proteins) quantitatively compared between untreated and ES2-treated neurons, where 88.5% of proteins remained unchanged with treatment. Significantly increased proteins are shown in red and significantly decreased protein shown in blue (p<0.05). C. Normalized intensity of biotinylated APP, APP-like proteins, and Notch-family proteins at the cell’s surface with ES2 as compared to untreated group. D. Normalized intensity of biotinylated APP-processing secretase proteins shows no significant changes after ES2 treatment for any secretases except for a slight increase of PSEN1. E. Normalized intensity of biotinylated proteins involved in neuronal transcytosis of APP revealed significantly differentially levels after ES2 treatment. Data were quality controlled, normalized, and statistically analyzed to identify differentially expressed features. Data are shown with ±SD and statistical significance was determined using unpaired Student’s t-test (*p < 0.05, **p < 0.01, ***p < 0.001, ns = not significant).

Multidimensional plot of all proteins captured through biotinylation show highest average intensity of canonical plasma membrane proteins (**Figure 1B**). Of the labeled cell-surface proteins, most of them remain unchanged with ES2 treatment (**Figure 1B**). There was a significant reduction of APP at the plasma membrane after ES2 treatment, but not APP homologs Amyloid Precursor-like Protein 1 and 2 (APLP1 and ALPL2) (**Figure 1C**). We also measured a reduction of Notch receptors after ES2 treatment (**Figure 1C**), which are similar type I transmembrane receptors processed by the same secretases and that functionally interact with APP ^35–38^. Interestingly, most of the secretase proteins involved in APP processing remained unchanged by ES2 treatment, except for an elevation of the active component of gamma-secretase (γ-Secretase), presenilin1 (PSEN1) (**Figure 1D**). Lastly, several endosomal proteins and motor proteins involved in APP transcytosis to the axon were detected by this biotinylation experiment and found to be reduced after ES2 treatment (Figure 1E). These identified proteins likely reflect the biotinylation and then subsequent internalization of cell-surface proteins, with streptavidin-resin capturing associated cytoplasmic proteins. This suggested that the exocyst may also be involved in APP trafficking through the endosomal pathway and transcytosis of APP C-terminal fragments through the axon ^39–41^.

### Exocyst inhibition results in reduced APP processing and amyloid-beta secretion

For a loss-of-function approach to further test if the exocyst has a role in the intracellular trafficking of APP and production of Aβ in neurons, we stably transfected SH-SY5Y cells with a human APP transgene containing familial AD Swedish (K595N/M596L) and Indiana (V717F) mutations, which are known to increase APP cleavage by BACE1 and thus Aβ production ^42^. Once this stable SH-SY5Y(mutAPP) transgenic cell line was established, we verified increased APP protein level and Aβ secretion into the cell medium by Western blot quantification and ELISA (**Figure 2A-C**). The SH-SY5Y(mutAPP) cell line secreted ∼550 pg/mL/24hrs of Aβ_40_, but only about ∼50 pg/mL/24hrs of Aβ_42_, which is consistent with physiological ratios of Aβ_40_/Aβ_42_. Due to the 10-fold higher Aβ40 secretion and sensitivity of ELISAs, we used Aβ_40_ levels in further experiments to assess changes of amyloidogenic processing due to exocyst disruption. We next generated a stable EXOC5 knockdown using shRNA in SH-SY5Y(mutAPP) cells, which yielded about a ∼70% decrease in EXOC5 protein levels (**Figure S2A,B**). Targeting the EXOC5 subunit has been successfully used in previous cell and animal models to disrupt the exocyst holocomplex function ^43^, since EXOC5 silencing significantly impacts stability of the holocomplex and other subunit functions in various tissues ^44–47^. In differentiated SH-SY5Y neurons, immunoblotting of cell lysates revealed an increase in full length APP after EXOC5 knockdown as compared to the SH-SY5Y(mutAPP) cells **(Figure 2A,D).** This indicates less APP is getting processed by α- and/or β-secretase with EXOC5 knockdown. Consistent with this finding, ELISAs revealed significantly decreased Aβ secretion by cells with knockdown EXOC5, which was comparable with decreases measured after treatment with γ-secretase inhibitor DAPT (**Figure 2E**).

**Figure 2:**
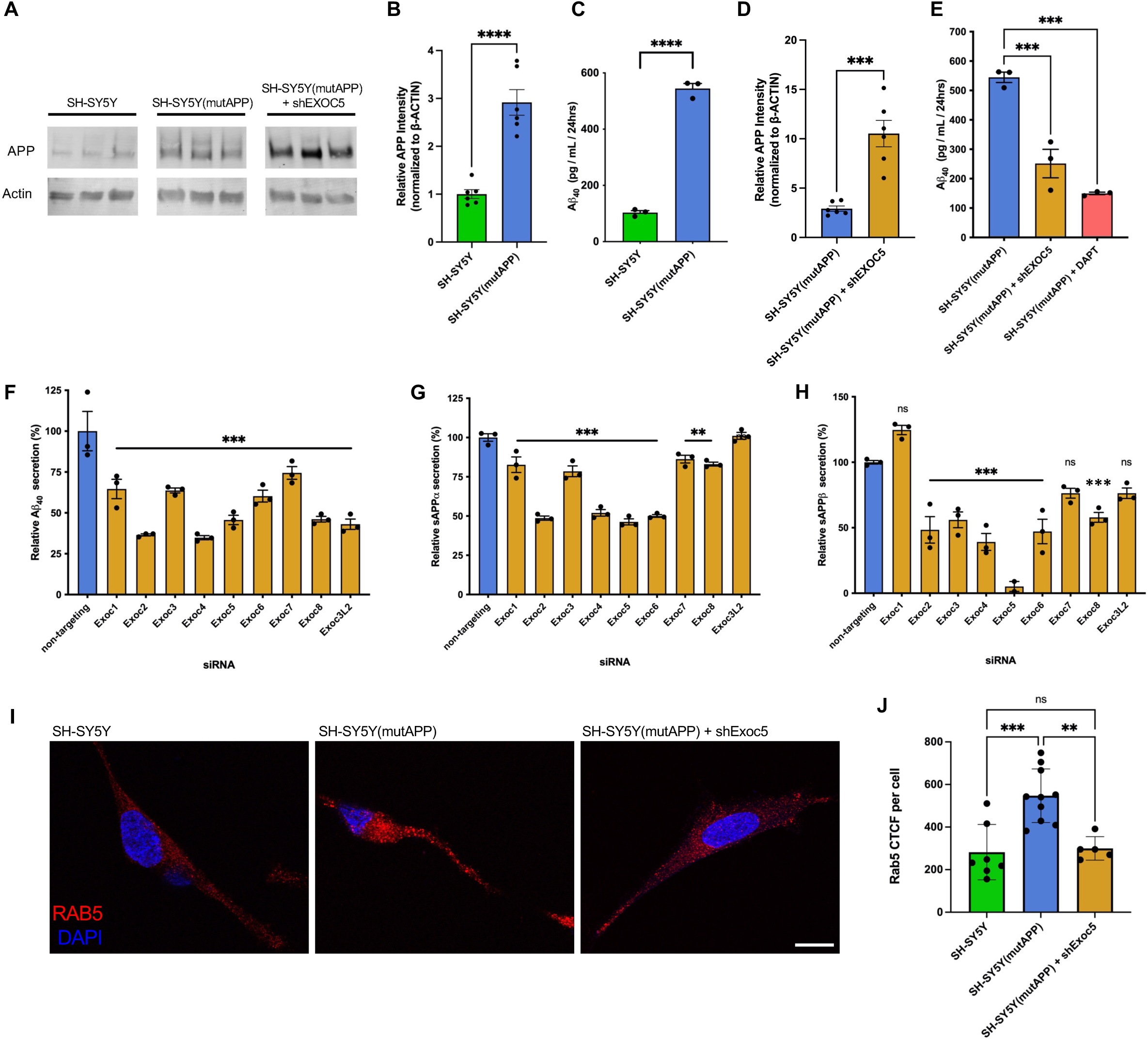
Exocyst inactivation in SH-SY5Y cells results in significant reduction of APP proteolytic processing and Aβ secretion. A. Representative immunoblotting of full length APP from cell lysates of our SH-SY5Y(mutAPP) cell line that expresses a APP transgene containing familial AD Swedish (K595N/M596L) and Indiana (V717F) mutations, showed an increase in full-length APP over the parental SH-SY5Y line. Also shown is further increased full length APP protein levels in SH-SY5Y(mutAPP) cells with shRNA-mediated stable knockdown of exocyst subunit EXOC5. Although cropped for space, all images are from a single immunoblot. B. Immunoblot quantification of full-length APP relative intensity normalized to β-actin confirmed higher APP protein levels in SH-SY5Y(mutAPP) cells than normal SH-SY5Y cells. C. ELISA measurements of conditioned cell medium revealed higher Aβ_40_ secretion by SH-SY5Y(mutAPP) as compared to normal SH-SY5Y cells. D. Immunoblot quantification of full-length APP relative intensity normalized to β-actin in SH-SY5Y(mutAPP) cells showed a significant increase after EXOC5 knockdown via stable expression of shRNA. E. ELISA measurements of SH-SY5Y(mutAPP) conditioned cell medium revealed a significant decrease in Aβ_40_ secretion after shRNA-silencing of EXOC5. Parallel controls included treatment of SH-SY5Y(mutAPP) cells with DAPT. F-H. ELISA measurements of SH-SY5Y(mutAPP) conditioned cell medium after siRNA knockdown of all eight individual exocyst subunits showed significantly decreased Aβ secretion and APP fragments (sAPPα and sAPPβ). I. Representative images from immunostaining of RAB5, an early endosome marker (red), in SH-SY5Y cells, SH-SY5Y(mutAPP), and SH-SY5Y(mutAPP) cells with shEXOC5 knockdown. Scale bar (white)= 10 µm J. Measurement of corrected total cell fluorescence (CTCF) for RAB5 immunostaining, showing a significant increase with mutAPP expression, which was reversed with shEXOC5 knockdown. Comparisons between two samples/conditions were performed using an unpaired Student’s t-test. Each datapoint for Western blots and ELISAs (A-H) represent a biological replicate, and each data point in RAB5 CTCF measurements (J) represent a field of view in a single experiment, representative of multiple independent experiments. Group values were compared with one-way ANOVA and corrected via post-hoc Tukey test. Significance value with ±SD shown as *(p<0.05), **(p<0.01), ***(p<0.001), **** (p<0.0001), and ns (not significant).

Moreover, we used small interfering RNAs (siRNAs) to individually knockdown all eight classic exocyst subunits (EXOC1-8). We transfected SH-SY5Y(mutAPP) cells with siRNAs against each of the exocyst genes, in parallel with a non-targeting control siRNA, replated the cells after 48 hours, and collected serum-free cell media and lysate after 72 hours. As measured by ELISA, knockdown of all eight exocyst subunits resulted in significantly decreased levels of secreted Aβ, compared to a non-targeting scrambled control siRNA (**Figure 2F, S2C**). We also used ELISAs to measure the soluble extracellular fragments of APP released into the media from these SH-SY5Y(mutAPP) cells after exocyst siRNA knockdowns, either from cleavage by α-secretase (sAPPα) or β-secretase (sAPPβ). Nearly all exocyst gene knockdowns yielded significant reductions of both sAPP*ɑ* and sAPPβ into the media (**Figure 2 G,H**).

Early endosomal swelling is commonly detected in neurons with increased rates of amyloidogenic APP processing, and can be detected by measuring levels of early endosomal protein markers, such as RAB5 ^39–41^. Confocal imaging of immunofluorescent-stained differentiated SH-SY5Y neurons confirmed a significant increase in RAB5 due to expression of the mutAPP transgene, but this was reversed after shRNA-mediated EXOC5 knockdown (**Figure 2 I,J**). Together, these data strongly supported the hypothesis that the exocyst complex has a critical role in regulating APP intracellular trafficking and proteolytic processing and is at least necessary for the initial delivery of APP to the somatodendritic plasma membrane. However, these results also allow for the possibility that the exocyst has other roles along the Aβ synthesis pathway.

### The exocyst and APP colocalize in neurons and display similar spatiotemporal dynamics, suggesting coordinated trafficking and mechanistic relationship in neuronal transport

Based on our previous loss-of-function experiments, we next assessed exocyst-APP colocalization and intracellular interaction by using immunofluorescence and immunofluorescent-tagged SH-SY5Y cell lines. Immunofluorescent staining visible through confocal microscopy revealed tight co-localization of EXOC5 and APP signals throughout primary mouse primary hippocampal neurons (pHCNs) (**Figure 3A,B**), and differentiated SH-SY5Y human neurons **(Figure 3C**). Overlap of EXOC5 and APP signals within the soma, and particularly along the neurites, suggest co-localization in the same subcellular regions of the neurons. To visualize overlapping APP and exocyst protein dynamics in living cells, three stable fluorescent SH-SY5Y neuronal cell lines were generated, each expressing a distinct exocyst subunit (Exoc1, Exoc4, or Exoc7) fused to a C-terminal mNeonGreen fluorescent tag (**Figure S2D**). These three subunits were chosen from a previously published analysis of which exocyst subunits could tolerate large epitope tags, such as fluorescent proteins, without disruption in their protein-protein interactions ^49^. The expression construct also included mutAPP fused to the red fluorescent protein mScarlet at the C-terminus. This enabled tracking of the full-length APP and also the internalized transmembrane APP CTFs, generated after α- or β-secretase cleavage, moving through the endosomal system and transcytosed into the axon. The expression and appropriate molecular weights of the tagged proteins were confirmed by Western blot analysis (**Figures S2E,F**).

**FIGURE 3:**
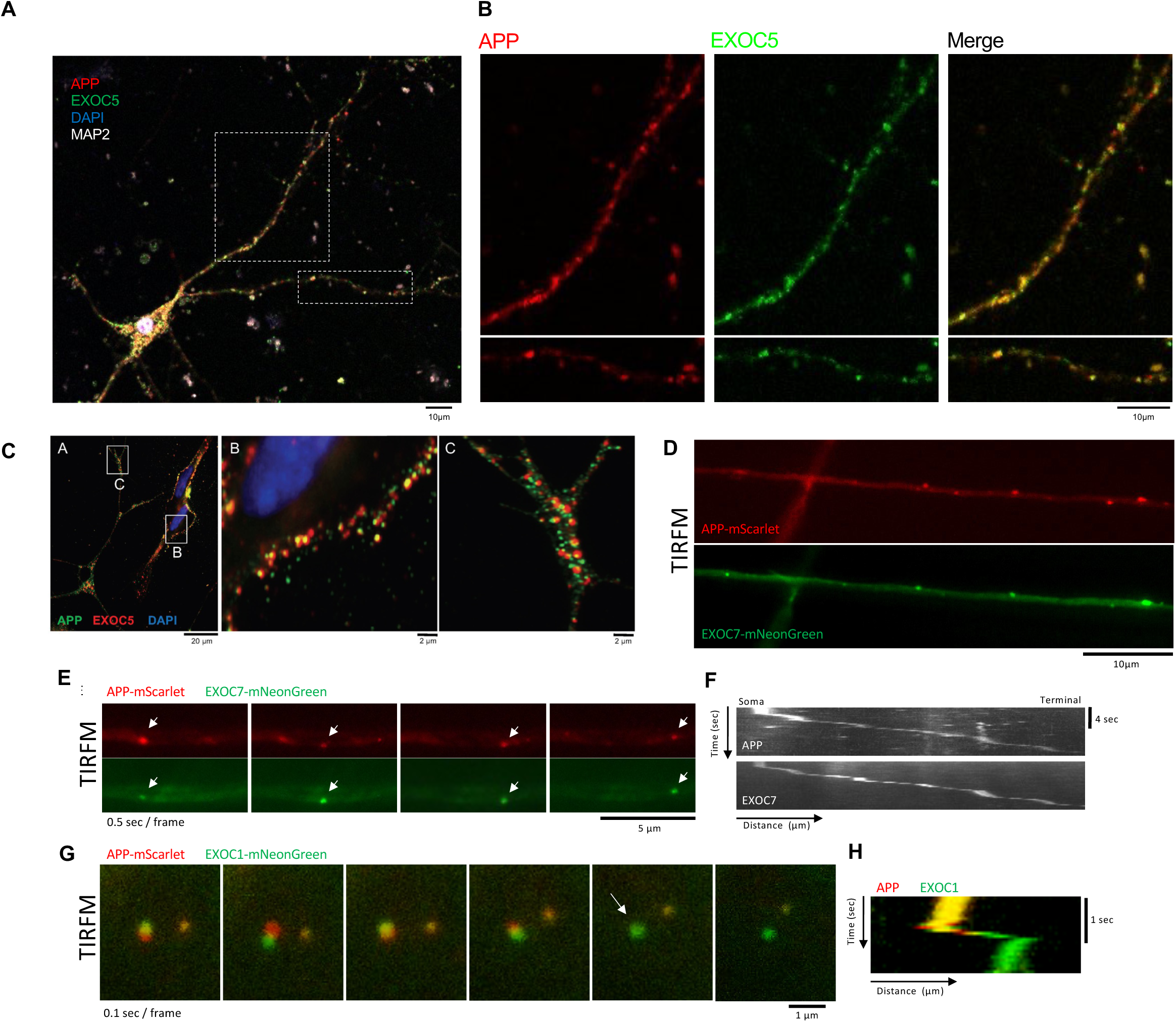
Exocyst proteins and APP co-localize at high resolution in mouse primary hippocampal neurons and differentiated SH-SY5Y neurons. A. Representative confocal images show a single mouse primary hippocampal neuron (pHCN) immunostained for EXOC5 (green), APP (red), and MAP2 (white). Nuclei were counterstained using DAPI (blue). B. Zoomed images from A with isolated color channels showing overlapping APP (red) and EXOC5 (green) puncta throughout the projections of mouse pHCN (merged=yellow). C. Representative confocal images show differentiated SH-SY5Y neurons immunostained for EXOC5 (red) and APP (green), with higher magnification of soma (b) and neurites (c). Nuclei were counterstained using DAPI (blue). D. Total internal fluorescence microscopy (TIRFM) images of living differentiated SH-SY5Y neurons co-expressing mutAPP-mScarlet and EXOC7-mNeonGreen fusion proteins demonstrated high-resolution colocalization in discrete particles along neurite projections. E. Time lapse, channel-split, images (0.5 sec/frame) from TIRFM of living differentiated SH-SY5Y neurons co-expressing mutAPP-mScarlet and EXOC7-mNeonGreen fusion proteins showed vesicles moving in a highly coordinated manner along a neurite. F. Kymographs of individual red (APP) and green (EXOC7) channels from the live-cell TIRFM video shown in E demonstrated overlapping APP/EXOC7 particle movement along SH-SY5Y neurites. Y-axis is time (sec) and X-axis is distance (µm). G. Time lapse images (0.1 sec/frame) from TIRFM of living differentiated SH-SY5Y neurons co-expressing mutAPP-mScarlet and EXOC1-mNeonGreen fusion proteins showed a punitive exocytosis event in the soma, observed in frame 5 (white arrow), with the disappearance of the APP signal (red) and remainder of EXOC1 signal (green). H. Kymograph of the live-cell vesicle shown in G highlighting colocalization of the two proteins and subsequent loss of APP signal (red). Y-axis is time (sec) and X-axis is distance (µm).

To visualize where APP and the exocyst may interact in living neurons, we performed live-cell TIRF microscopy of differentiated SH-SY5Y neurons co-expressing these APP-mScarlet and exocyst-mNeonGreen fusion proteins. These experiments revealed high-resolution colocalization between APP and each of the fluorescently labeled exocyst subunits, particularly in discrete particles moving along neurite projections (**Figure 3D**). A representative time-lapse sequence (0.5 sec/frame) showing highly coordinated movement between APP-mScarlet and EXOC7-mNeonGreen is depicted in **Figure 3E**, with the associated kymograph shown in **Figure 3F**. A putative exocytosis event is shown in **Figure 3G**, further illustrated by the corresponding kymograph (**Figure 3J**) showing the transient association of the two proteins, followed by a loss of APP-mScarlet signal that could be consistent with membrane fusion and release. Large puncta distributions from labeled exocyst subunits are likely from assembly/clustering of multiple exocyst octamers at various exocytic docking sites ^49–50^. These high-resolution imaging approaches support the hypothesis that the exocyst complex likely plays a critical role in the intracellular trafficking of APP-containing vesicles throughout various regions of neuronal cells. These results further compliment loss of function experiments where APP trafficking and Aβ secretion is significantly disrupted if exocyst is inhibited.

### Chemical inhibition of exocyst leads to reduced APP trafficking into the axon

After establishing that APP localization to the plasma membrane was significantly decreased by ES2 treatment in our differentiated 5Y cells and that exocyst knockdown decreased Aβ generation measured in SH-SY5Y(mutAPP) cells, we wanted to test whether axonal trafficking of APP+ vesicles was also affected by exocyst inhibition. To do this, we utilized mouse pHCN and treated them with ES2 (or vehicle) as previously described, then fixed them for immunostaining and confocal microscopy. Based on previous findings regarding Aβ release at the pre-synapses and APP-CTF vesicles translocation into the axon ^51–53^, we measured APP+ vesicles in the axon initial segment (AIS) with and without ES2. The AIS is a specific neuronal region that demarcate the separation of the soma from the axon proper and is structured by various proteins such as AnkyrinG (ANKG) (**Figure 4A**) ^54–55^. Pearson’s correlation coefficient (PCC) analysis via FIJI software was used to analyze correlation of APP with ANKG (**Figure 4B**), and also the EXOC5-ANKG correlation and the APP-EXOC5 correlation specifically in the AIS segment (**Figure 4C**). We found that ES2 treatment significantly decreased APP-ANKG correlation (**Figure 4D**), suggesting less APP is trafficked to the AIS when the exocyst is inhibited. No significant difference was detected with EXOC5-ANKG correlation with ES2 treatment (**Figure 4E**) implying that exocyst proteins are still within the AIS. However, with less APP detected in the AIS, EXOC5-APP colocalization within the AIS was also decreased (**Figure 4F**). We also wanted to know whether the detected decrease in EXOC5-APP colocalization with ES2 treatment was uniform across the entire length of the AIS and whether ES2 treatment could potentially affect AIS lengths. Using a scatterplot comparison of EXOC5-APP PCC against AIS length, we didn’t detect significant correlation (**Figure 4G**). This highlights that under ES2 treatment, the decrease in EXOC-APP colocalization was detected evenly throughout the entire length of the AIS, as opposed to a specific subsection, without altering the AIS scaffold itself. Overall, these findings show that chemical inhibition of the exocyst results in less APP+ vesicles trafficked into the axon, as well as significant decreases in co-localized exocyst and APP in the AIS segment.

**FIGURE 4:**
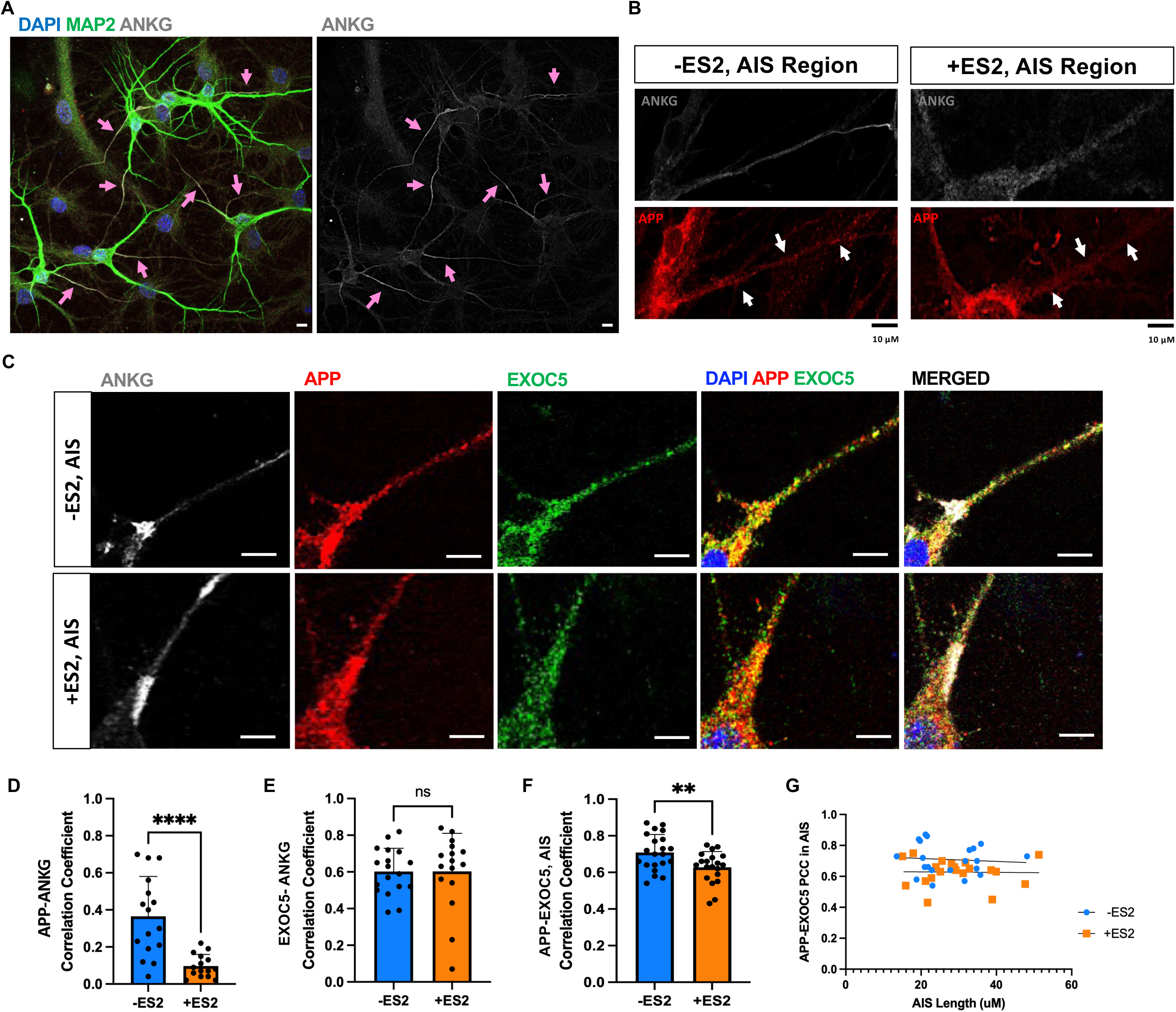
Chemically inhibiting exocyst disrupts intracellular APP trafficking into the axon initiation segment (AIS) of mouse primary hippocampal neurons, identified by Ankryin G immunostaining (ANKG). A. Representative immunostaining and confocal microscopy of mouse primary hippocampal neurons of the axonal initial segment at baseline and with the exocyst inhibitor, endosidin2 (ES2) denoted by ANKG (gray) and pink arrows. Nucleus stained by DAPI, EXOC5 (red) and dendrites with MAP2 (green). White scale bar= 20 µm B. Representative close-up confocal imaging of ANKG (gray) and APP (red) immunostaining in mouse primary hippocampal neurons with and without ES2 treatment. White arrows show APP staining throughout the AIS region. Black scale bar= 10 µm C. Representative immunostaining and confocal microscopy of ANKG (gray), APP (red), EXOC5 (green), with the merged image on the right. Nucleus represented by DAPI staining. White scale bar: 5 µm D. Pearson’s correlation coefficient (PCC) measuring the correlation between APP and AnkG stained regions showed significant decrease in APP correlation with ANKG E. PCC of EXOC5-ANKG colocalization showed no significant changes after ES2 treatment. F. PCC analysis of colocalized EXOC5-APP within the AIS showed significant decrease after ES2 treatment. G. Scatterplot plotting APP-EXOC5 PCC against AIS length (µm) showing no correlation along the entire length of the axon suggesting PCC is consistent throughout the AIS and the treatment did not shorten or lengthen the AIS region. Statistical comparison between two samples/conditions was performed using an unpaired Student’s t-test. Each dot in D-F represents the PCC measured from the AIS of an individual neurons pooled across 3 independent coverslips, sampled from a minimum of 6 fields of view. Significance value with ±SD shown as *(p<0.05), **(p<0.01), ***(p<0.001), **** (p<0.0001), and ns (not significant). Scale bar (white)= 5 µm.

### Exocyst colocalizes with APP and GLUT4 containing vesicles in the hippocampal region of mouse brains

To visualize exocyst-APP associations within hippocampal regions, proximity ligation assays (PLA) were performed on wild type coronal mouse brain sections. Hippocampal region signals were quantified via FIJI object counter and compared to the thalamic region and normalized against number of nuclei per microscopic field, showing a significant increase of EXOC5-APP PLA signal specifically in the hippocampus as compared to the thalamic region (**Figure 5A,B**). These results validate the immunocytochemistry colocalization and correlation results previously observed in SH-SY5Y neurons and pHCNs and provides larger functional context in the hippocampus as an early site of Aβ accumulation ^58–59^.

**FIGURE 5:**
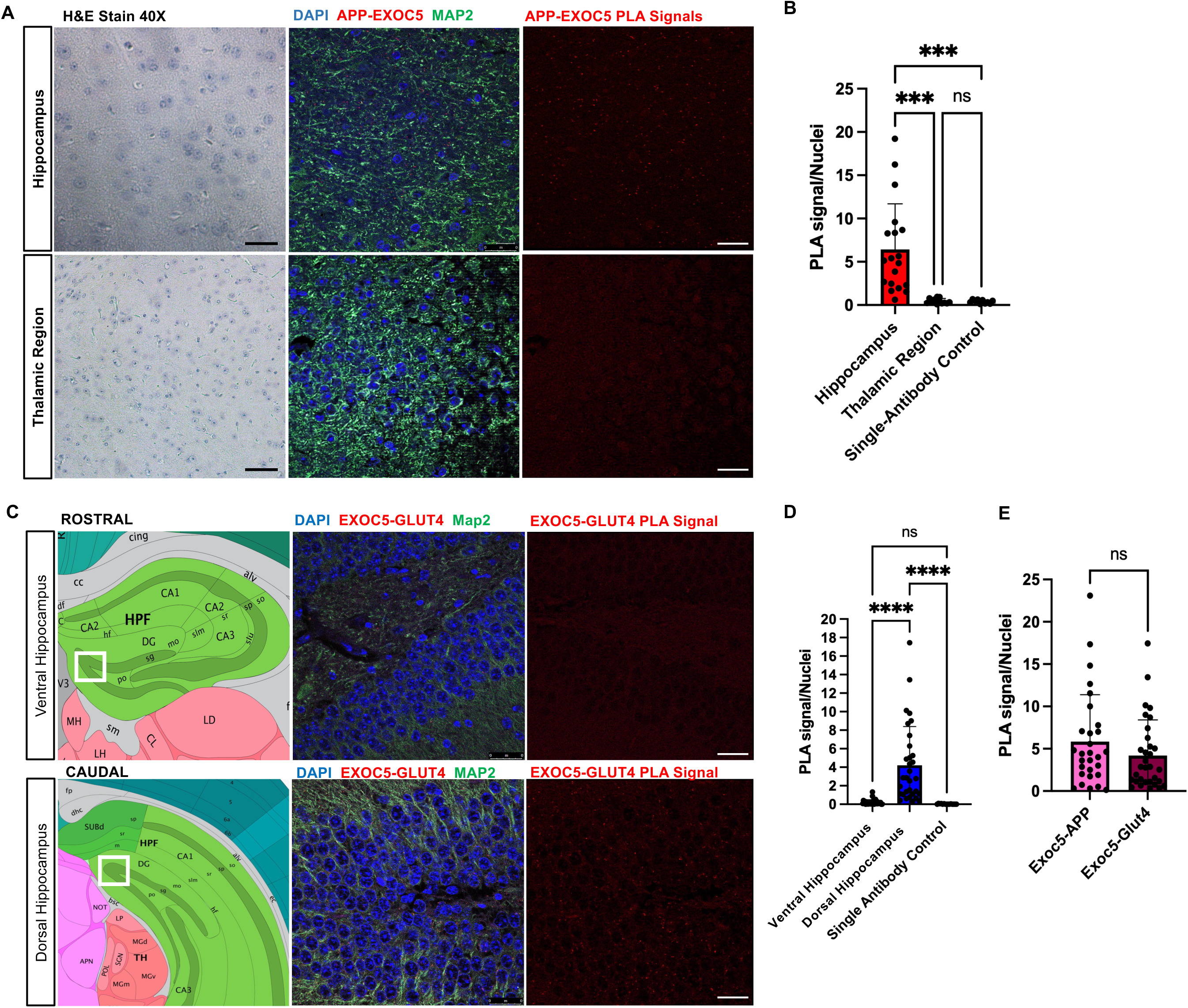
Proximity ligation assays show EXOC5-APP and EXOC5-GLUT4 associations in hippocampus of wild type mouse brains. A. Representative images of coronal mouse hippocampal and thalamic brain regions probed for APP-EXOC5 associations (PLA signal observed in red) along with hematoxylin & eosin staining (40X) for visual reference. Dendrites are marked by MAP2 immunostaining (green) and nuclei with DAPI (blue). B. Quantification of PLA signals showed APP-EXOC5 associations (red puncta) are significantly higher in hippocampal regions compared to the thalamic regions. Single-antibody negative controls were also run in parallel and showed very few to no PLA signal. C. Schematic images of the coronal mouse hippocampal regions of PLA analysis (white box) from the Allen Mouse Brain Atlas along with representative PLA images of coronal mouse hippocampal and thalamic brain regions probing for EXOC5-GLUT4 associations (red puncta) along with nuclei marked by DAPI (blue) and dendrites by MAP2 (green) immunostaining. D. Quantification of PLA signals showed EXOC5-GLUT4 associations (red) within the ventral and dorsal coronal hippocampal slices, with a significant increase of EXOC5-GLUT4 PLA signal in the dorsal hippocampal region. E. Quantification of PLA signals showed similar EXOC5-APP and EXOC5-GLUT4 distributions within coronal hippocampal slices, with no significant differences. Group comparisons utilize one-way ANOVA with post-hoc Tukey test. Each condition represents n=3 technical replicates from 3 biological replicates of coronal mouse hippocampal slices. Significance value with ±SD shown as *(p<0.05), **(p<0.01), ***(p<0.001), **** (p<0.0001), and ns (not significant). Data points represent at least 3 microscopic fields of view within the highlighted regions (white box), Black scale bar: 40 µm, white scale bar: 25 µm.

Given the critical role of the exocyst for the exocytosis of GLUT4 after insulin signaling in peripheral tissues, we wanted to characterize the baseline distribution of EXOC5-GLUT4 associations in the hippocampus to see if this would overlap with EXOC5-APP signal. The EXOC5-GLUT4 PLA did reveal significantly more numerous puncta per nuclei as compared to the single-antibody control. In contrast to the EXOC5-APP, which was consistent through the hippocampus, the distribution of EXOC5-GLUT4 signal throughout ventral and dorsal hippocampal regions were highly variable with ventral slices having significantly less EXOC5-GLUT4 PLA signals as compared to the dorsal slices (**Figure 5C,D**). This may be reflective of regional differences in the hippocampus when comparing the dentate gyrus to the CA1-3 regions based on increased neuronal activity ^60–62^. Lastly, when comparing the level of EXOC5-APP PLA signals to EXOC5-GLUT4 PLA signals in the dorsal hippocampus, there was no significant differences (**Figure 5E)**. This may indicate that the exocyst interacts with both APP and GLUT4 in shared exocyst-mediated trafficking pathways not only in cultured cell populations, but at the hippocampal tissue level.

### Insulin signaling directs exocyst activity away from APP+ vesicles and towards GLUT4+ vesicles

In peripheral metabolic tissues, such as adipocytes and skeletal muscle, insulin signaling plays a key role in regulating exocyst assembly, vesicle targeting, and activity ^22–23,63^. The most well-characterized pathway is the rapid stimulation by insulin of exocyst-dependent GLUT4 exocytosis onto the plasma membrane to boost glucose uptake. However, whether this pathway is conserved in neurons has not been previously investigated. To test if the exocyst indeed has a role in regulating GLUT4 exocytosis in neurons, we cloned a human GLUT4 transgene containing an mScarlet fluorescent protein on the C-terminus, and a pHluorin fluorescent protein inserted in-frame in the first exofacial loop at amino acid 67 (**Figure 6A**). Placement of the pHluorin tag at this particular location has been shown to allow for proper GLUT4 trafficking and channel activity, including in neurons ^67–68^. The pHluorin is a pH-sensitive GFP fluorophore which has low fluorescence in low pH environments (such as intracellular transport vesicles and endosomes), but high fluorescence in the neutral pH of the cell culture media ^69–72^. Thus, measuring green fluorescence correlates with the amount of GLUT4 protein on the plasma membrane surface, and can be normalized for total GLUT4 with red fluorescence. SH-SY5Y cells were transfected with our pGENIE3-pHluorin-GLUT4-mScarlet expression plasmid and were plated in 96-well plates, fully differentiated into neurons, and green and red fluorescence were read using a microplate reader. Under baseline conditions, both green and red fluorescence levels were correlated with seeded cell number (**Figure S3A,B**), and by normalizing pHluorin/mScarlet signal, we were able to measure a strong drop in green fluorescence (but not red) when cells were incubated in acidic cell medium (pH 5) (**Figure S6B**). We also detected a steady decrease of pHluorin signal with insulin withdraw over time (**Figure S6C**), and sharp increase after only 5 minutes of insulin addback (**Figure S6D**). Using ES2 to inhibit exocyst activity, we found that ES2 treatment steadily lowers pHluorin signals in these cells under basal serum conditions, but not mScarlet signals (**Figure 6E,F**). Moreover, similar to the PI3K inhibitor, LY294002, ES2 strongly prevented the rapid increase of pHluorin signaling induced by 5 minutes of insulin addback (**Figure 6G**). Altogether, these results suggest that the exocyst is necessary for insulin-driven GLUT4 exocytosis in neurons, and this pathway that has been well described in peripheral metabolic tissues is conserved in the central nervous system. However, in this context, GLUT4 trafficking likely supports rapid energy utilization associated with memory processes rather than long-term metabolic homeostasis ^64–66^.

**FIGURE 6:**
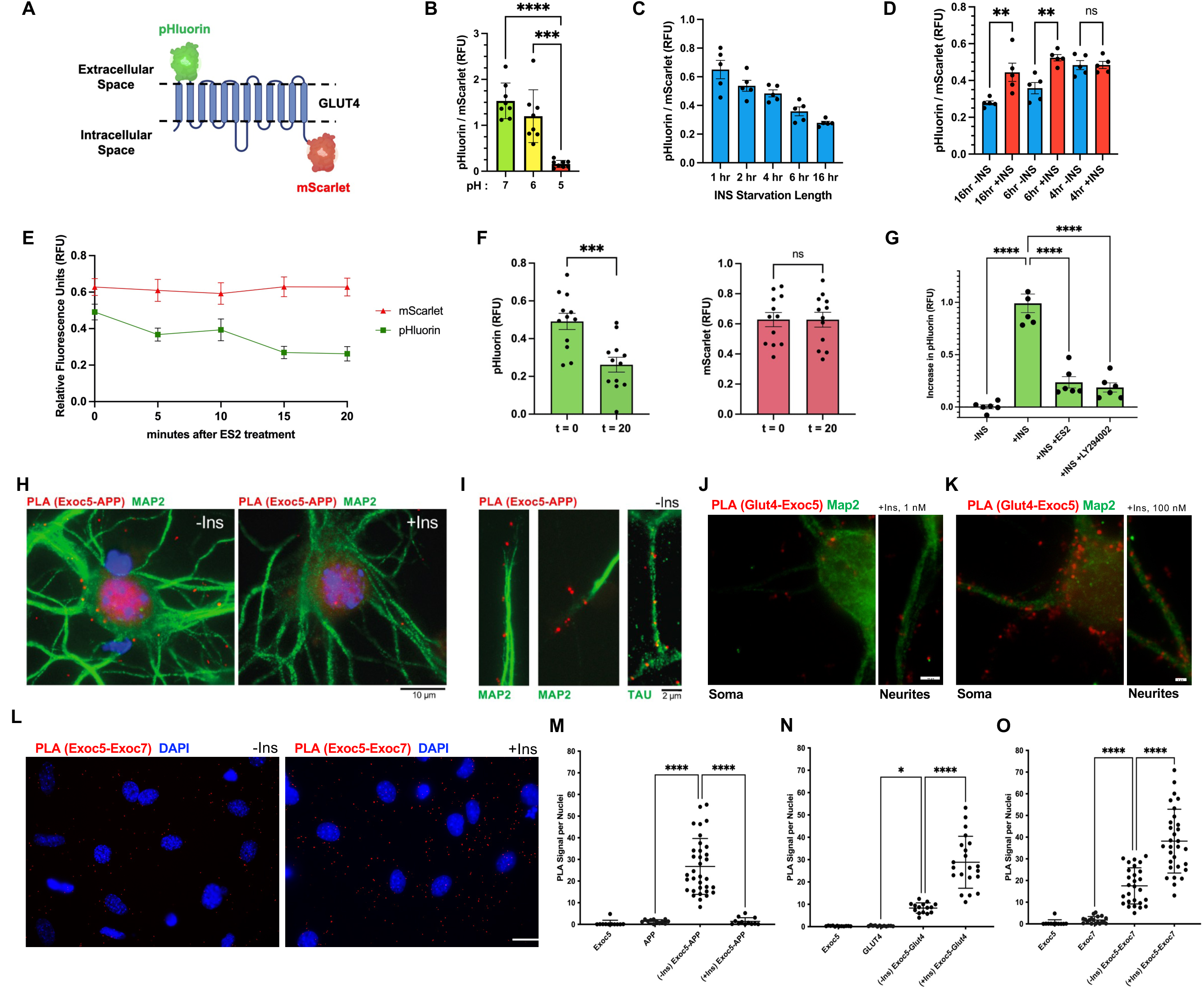
Insulin signaling in neurons redirects the exocyst complex away from APP+ vesicles towards GLUT4+ vesicles for GLUT4 exocytosis. A. Schematic representation of the pHluorin-GLUT4-mScartlet fluorescent fusion protein stably expressed in SH-SY5Y cells. The modified GFP pHluorin on the first exofacial loop remains dim in acidic endosomal or TGN compartments but fluoresces strongly when GLUT4 is exocytosed to the plasma membrane and pHluorin is in a neutral pH environment. The mScarlet RFP attached to the C-terminus of GLUT4 provides constant red fluorescence as an internal control. B. Fluorescent microplate reading of differentiated SH-SY5Y neurons stably expressing pHluorin-GLUT4-mScarlet showed pHluorin signal decreased when the pH of the cell medium was lowered, confirming pH sensitivity of pHluorin. C. Fluorescent microplate reading of differentiated SH-SY5Y neurons stably expressing pHluorin-GLUT4-mScarlet showed decreasing pHluorin/mScarlet signal with longer insulin starvation times. D. Comparison of before and after 15 minutes of 100nM insulin addback after varying lengths of insulin starvation. Longer starvation periods yielded greater significant increases from insulation stimulation. Statistical analysis was completed using a paired T-Test as the same biological replicates during starved conditions were measured after insulin treatment. E. Microplate fluorescence reading of differentiated SH-SY5Y neurons expressing pHluorin-GLUT4-mScarlet under basal culture conditions showed a steady reduction in pHluorin GFP fluorescence (green) with ES2 incubation, while mScarlet (red) remained unchanged. F. Quantitative analysis from data shown in J at 0 minutes as compared to 20 minutes of ES2 treatment revealed significant pHluorin fluorescence decreases but unchanged mScarlet signal. G. Microplate fluorescence reading of the change in pHluorin fluorescence signal in differentiated SH-SY5Y neurons after an overnight insulin starvation followed by 5 minutes of a 100 nM insulin addback (+INS). This increase in pHluorin signal was significantly prevented by treatment with ES2, or the PI3K inhibitor (LY294002), and absent with no insulin addback (-INS). H-K. Representative images of PLA signals (red puncta) with immunofluorescent counterstaining in isolated mouse pHCN, under insulin starved and insulin addback conditions. Robust EXOC5-APP association was observed during insulin starvation conditions (-Ins), with rapid PLA reduction within minutes of insulin addback (+Ins). Conversely, APP-GLUT4 associations were noticeably increased with insulin treatment. Nuclei stained by DAPI (blue), dendrites by MAP2 and axons by TAU immunostaining (both green). L. Representative images for quantitative PLA analysis of exocyst associations in pHCNs, under insulin-starved (-Ins) and after 15 of 100 nM insulin addback (+Ins) conditions, with DAPI counterstain. Images show EXOC5-EXOC7 PLAs to reflect assembly of the exocyst holocomplex. PLA signal in each microscope field is normalized by cell nuclei. F. Quantification of EXOC5-APP PLA signal in pHCNs after 2 hours of insulin starvation, and after 15 minutes of 100 nM insulin addback, showed a dramatic decrease in EXOC5-APP association after insulin treatment. Parallel PLAs with single antibodies were performed as negative controls. G. Quantification of EXOC5-GLUT4 PLA signal in pHCN under the same conditions conversely showed a significance increase in EXOC5-GLUT4 association after insulin treatment. H. Quantification of EXOC5-EXOC7 PLA signal in pHCN under the same conditions showed an overall increase in EXOC5-EXOC7 association with insulin treatment. Group analyses were completed with one-way ANOVA and post-hoc Tukey’s test while comparison within two groups were completed using unpaired Student’s t-test. Each florescence data point (B-G) represents a biological replicate (microplate well). Each PLA data point (M-O) represents a field of view from an experiment with pooled pHCN isolation, representative of three independent trials. Significance value with ±SD shown as *(p<0.05), **(p<0.01), ***(p<0.001), **** (p<0.0001), and ns (not significant). Scale bar values are noted within figure.

We next utilized PLA to compare exocyst assembly and targeting to APP+ or GLUT4+ vesicles, under insulin-starved or insulin-stimulated conditions, in both differentiated SH-SY5Y cells and mouse pHCN cultures. Cells were grown on glass coverslips, starved in insulin-free medium for 1 hour and then treated with 100 nM insulin, or left untreated in insulin-starved conditions. Cells were fixed and PLA was performed with antibodies against EXOC5 and APP to measure associations between the two proteins. In pHCNs, substantial decreases in EXOC5-APP PLA signals were measured after insulin treatment while PLA signals were significantly increased during insulin withdrawal with distributions throughout the soma and along dendrites and axons marked by MAP2 and TAU (**Figure 6H-I**) and quantified as puncta per nuclei (**Figure 6M**). Simultaneously, EXOC5-GLUT4 PLA signals were few during insulin withdrawal in the soma and neurites and significantly increased after insulin treatment (**Figure 6J-K**) validated through quantification as well (**Figure 6N**). Representative images from PLAs of exocyst subunits (EXOC5-EXOC7) show that the exocyst holocomplex assembly is present during insulin withdrawal but is significantly elevated after insulin addition (**Figure 6L,O**). We found the same changes in exocyst assembly and contrasting associations with APP or GLUT4 with PLAs in differentiated SH-SY5Y neurons under insulin starvation and after insulin addback (**Figure S3C**). Collectively, these findings suggest that the exocyst has an inverse relationship between trafficking APP+ vesicles and GLUT4+ vesicles, and insulin signaling plays a key role in this balance.

## DISCUSSION

In this study, we identify the exocyst trafficking complex as a previously unrecognized regulator of amyloid precursor protein (APP) intracellular trafficking and proteolytic processing in neurons. Using complementary proteomic, genetic, imaging, and functional approaches, our findings demonstrate that disruption of exocyst activity reduces APP delivery to the neuronal plasma membrane, alters axonal sorting, and significantly decreases amyloidogenic processing and amyloid beta (Aβ) secretion. Importantly, we further show that insulin signaling dynamically regulates this pathway by redirecting exocyst activity away from APP-containing vesicles and toward GLUT4 trafficking. Together, these results establish a mechanistic framework linking vesicle tethering machinery, neuronal insulin signaling, and APP processing pathways relevant to Alzheimer’s disease pathogenesis.

Intracellular trafficking plays a central role in determining whether APP undergoes amyloidogenic or non-amyloidogenic processing. Previous studies have emphasized the importance of endosomal localization and axonal transport in regulating access of APP and its C-terminal fragments to secretase complexes ^1–3, 73–75^. Our data suggest that the exocyst is central to these activities by regulating delivery of APP-containing vesicles to the somatodendritic plasma membrane and subsequent routing toward axonal compartments. Acute pharmacologic inhibition or genetic disruption of exocyst components reduced surface APP abundance and resulted in accumulation of full-length APP with concurrent reductions in both soluble APP fragments and secreted Aβ. These findings support a model in which exocyst-dependent vesicle targeting is required for efficient entry of APP into productive processing pathways rather than directly regulating secretase activity itself.

Several observations further support a coordinated trafficking relationship between APP and the exocyst complex. Live-cell imaging demonstrated dynamic colocalization between APP and multiple exocyst subunits during vesicle movement and putative exocytic events, while chemical inhibition altered APP localization within the axon initial segment without broadly disrupting exocyst localization. These results suggest cargo-selective regulation rather than generalized trafficking failure. The reduced delivery of APP to the axon initial segment following exocyst inhibition is particularly notable given increasing evidence that final γ-secretase processing occurs predominantly at presynaptic regions ^76^. Altered routing at this early axonal checkpoint may therefore substantially influence downstream Aβ production. Consistent with this interpretation, exocyst disruption also reversed Rab5-associated early endosomal enlargement observed in APP-overexpressing neurons, a phenotype strongly associated with enhanced amyloidogenic processing in Alzheimer’s disease models ^77^.

Although the exocyst has well-established roles during neuronal development ^15,18,22,78–79^, its function in mature neurons has remained unclear. Our findings suggest that one important role of the complex may be selective regulation of long-range membrane cargo trafficking rather than synaptic vesicle release. Notably, several small GTPases previously implicated in exocyst regulation, including Rab and Arf family members, have also been linked independently to APP trafficking and Alzheimer’s disease risk pathways ^80–82^. The present results place the exocyst as a potential organizing complex integrating these trafficking signals within neurons.

A major conceptual advance from this work is the identification of insulin signaling as an upstream regulator of exocyst-dependent APP trafficking. Epidemiologic and clinical studies have long associated metabolic dysfunction and brain insulin resistance with increased Alzheimer’s disease risk ^83–86^, yet the molecular mechanisms underlying this relationship remain incompletely defined. Here, insulin stimulation promoted exocyst assembly while simultaneously decreasing exocyst association with APP and increasing interactions with GLUT4-containing vesicles. Functional assays confirmed that neuronal GLUT4 exocytosis requires exocyst activity and is rapidly stimulated by insulin signaling. In order to measure changes in trafficking and exocytosis in the timeframe of minutes, we used 100 µM of insulin to rapidly saturate receptors and induce a robust synchronized signal. However, this high nanomolar insulin concentration may potential also activate IGF-1R, so further work may have to dissect the insulin signaling pathways that regulate the exocyst in neurons. These findings support a competitive or cargo-selective trafficking model in which metabolic signaling directs exocyst resources toward glucose uptake pathways at the expense of APP transport. Under conditions of impaired insulin signaling, increased association between APP and the exocyst may favor trafficking routes that enhance amyloidogenic processing. While our data demonstrates that insulin bioavailability modulates exocyst associations with APP+ and GLUT4+ vesicles in neurons, we acknowledge this may reflect a complex modulation of intracellular trafficking of neuronal vesicles as a whole rather than only a specific redirection of exocyst.

The regional enrichment of exocyst interactions with both APP and GLUT4 within the hippocampus further supports potential disease relevance, as this region is particularly vulnerable during early Alzheimer’s disease progression ^80, 87^. The ability of insulin signaling to dynamically redistribute trafficking machinery may therefore represent an important mechanism linking neuronal metabolic state to amyloid production in selectively vulnerable neuronal circuits.

As a limitation of this study, much of the mechanistic analysis was performed in differentiated neuronal cell systems, and future studies will be required to determine how exocyst-dependent APP trafficking operates *in vivo* during aging or under metabolic stress. In addition, while our findings demonstrate a requirement for the exocyst in APP processing, the precise molecular determinants governing cargo selection between APP and GLUT4 vesicles remain unknown. Defining how insulin signaling pathways regulate exocyst targeting in neurons will be an important area for future investigation.

In summary, our results support a model in which the exocyst complex regulates neuronal APP trafficking and amyloidogenic processing while serving as a molecular intersection between vesicle transport and insulin signaling pathways. These findings provide new insight into how metabolic dysfunction may influence Alzheimer’s disease risk through regulation of intracellular trafficking mechanisms rather than direct modulation of secretase activity alone.

## METHODS

### Animals

All animal procedures and protocols were approved by the University of Hawaii IACUC (protocol #11-1094), under the University of Hawaii‘s Animal Welfare Assurance (A3423-01) on file with the Office of Laboratory Animal Welfare. Mice were housed under standard conditions with a 12-hour light cycle with water and food *ad libitum*.

### Cell Cultures and Reagents

SH-SY5Y cells (Sigma-Aldrich) were cultured following recommendations by American Type Culture Collection using Dulbecco’s Modification of Eagle’s Medium (DMEM)/Ham’s F-12 50/50 mix media with 10% fetal bovine serum (FBS) and penicillin-streptomycin. SH-SY5Y cells were differentiated into neurons following the 18-day protocol described in Shipley et al.^46^ Briefly, the day after seeding, cells were cultured in Medium 1 (EMEM, 2.5% FBS, 1x pen-strep, 2mM glutamine, 10 μm retinoic acid (RA)) from Days 1-7, then Medium 2 (EMEM, 1% FBS, 1x pen-strep, 2 mM glutamine, 10 μm RA) from Day 8, and then Medium 3 (Neurobasal media, 1x B27, 1x N2, 1x pen-strep, 2mM glutamine, 50 ng/mL BDNF, 10 μm RA) from Day 10-18.

Mouse primary neurons were isolated from the hippocampus removed from neonatal (P0-3) C57Bl6/J wild type (WT) mice and cultured to 12 days *in vitro* (DIV12). Hippocampi were isolated, pooled from littermates, and digested in papain solution containing 10mM L-cysteine and cell dissociation buffer, and then incubated at 37°C for 15 min. After recovery by centrifugation, the tissue was suspended in Neurobasal A (w/FBS) and dissociated by sequential trituration. Cells were pre-plated to remove adherent non-neuronal cells (primarily glia), and the neuron-enriched suspensions were plated onto poly-D-lysine or CellTak (Corning) coated coverslips and cultured in Neurobasal A medium containing 5% FBS, B-27 supplement, and gentamicin.

In some experiments, 100 µM of endosidin-2 (ES2) was used to inhibit the exocyst, based on previously published analyses of effective dosages in mammalian cells ^30–33^. For RNAi knockdowns, SH-SY5Y cells were seeded at 70% confluency and transfected with 50ng of siRNA (Dharmacon SMARTpools) or EXOC5 shRNA (Sigma) using Lipofectamine 3000 Reagent (Thermo Fisher). All transgene expression plasmids were generated using the pGENIE3 (pG3)-piggyBAC transposon system backbone ^90^, which has hygromycin resistance for mammalian cell selection. Plasmids were cloned in-house, sequence-verified, and transfected into undifferentiated SH-SY5Y cells via Lipofectamine 3000. Proper expression of proteins-of-interest were verified using western blotting, as previously described ^91^. Stable transgenic SH-SY5Y cell lines were generated for each plasmid through continuous hygromycin selection.

For insulin starvation experiments, fully differentiated neurons were incubated with either Medium 3 (for SH-SY5Y neurons) or serum-free Neurobasal A with Glutamax (for primary neurons), but each supplemented with insulin-free B27 (Thermo Fisher #A1895601) for shown lengths of time. For insulin addback experiments, 100 nM insulin (Gibco) in the appropriate medium above was incubated with the neurons for 5-15 minutes after a period of insulin starvation. In some experiments, controls included the PI3K inhibitor LY294002 (10 µM, Calbiochem), and the γ-secretase DAPT (10 µM, Cayman Chemical).

### Biotinylation of Cell Surface Proteins and Proteomics

Fully differentiated SH-SY5Y cells in Medium 3, either treated for 2 hours with 100 µM ES2 or vehicle control, were washed with 1X PBS and surface proteins underwent biotinylation for 10 minutes at RT using the Pierce Cell Surface Biotinylation kit (#A44390). The cells were then lysed and biotinylated proteins were captured on neutravidin resin columns provided in the kit. After extensive washing, column-bound proteins were eluted with dithiothreitol (DTT) and purified eluates were precipitated using an acetone precipitation method, resuspended in RIPA buffer, and frozen at -80°C. Protein digestion and mass spectrometry analysis were performed at the IDeA National Resource for Quantitative Proteomics at the University of Arkansas for Medical Sciences (UAMS), as fully described in Supporting Information (SI).

### Enzyme-linked Immunosorbent Assay (ELISA)

To measure the rate of secreted Aβ, conditioned serum-free Neurobasal cell medium was collected from SH-SY5Y cells cultured in microplates. High Sensitivity Human Amyloid β ELISA kits (Millipore) were used to measure either Aβ_40_ (#EZHS40) or Aβ_42_ (#EZHS42). Standard curves were generated using the kits’ positive control peptides, and quantities of Aβ in 200 µL of media samples were interpolated from the standard curves.

### Immunocytochemistry and Proximity Ligation Assay (PLA)

Neurons cultured and differentiated on glass coverslips were washed three times with PBS and fixed for 10 minutes with 4% PFA. Cells were permeabilized for 10 minutes with 0.1% Triton X-100, blocked in 0.1% BSA in PBS, incubated with primary antibodies overnight at 4°C, incubated with secondary antibodies for 1 hour, washed, and mounted using VECTASHIELD® Antifade Mounting Medium. See Supplementary Table 1 in SI for relevant antibodies and combinations. For PLA on SH-SY5Y neurons and pHCNs, cells were seeded and differentiated on glass coverslips before being fixed for 10 min in 4% PFA, permeabilized in 0.1% Triton-X 100 in PBS for 10 min, and stored in 0.1% BSA. For PLA on mouse brain histological sections, antigen retrieval was performed as detailed in SI. PLA was performed according to manufacturer instructions using the DuolinkTM In-Situ Red Starter Kit (Mouse/Rabbit) (Millipore). Neurons or tissues sections were incubated overnight at 4°C with the antibody combinations EXOC5-EXOC7, EXOC5-GLUT4, or EXOC5-APP, with additional single antibody negative controls. Samples were washed and incubated for 1 hour at 37°C in a humidity chamber with the PLUS and MINUS antibody probes. Samples were washed again and then incubated with ligase at 37°C in a humidity chamber for 30 min. Lastly, samples were washed and incubated with polymerase and appropriate buffer for 100 min at 37°C in a humidity chamber. Co-staining was done using anti-MAP2 or anti-TAU with Alexa-488 secondary antibodies or using Alexa Fluor 488 phalloidin (Thermo Fisher) to stain F-actin. PLA signal and counterstains were imaged with an Olympus BX41 fluorescent microscope. Quantification was done by taking 10 to 40 images of each condition, with each frame containing 5-15 cells. Images were analyzed with ImageJ software, using the detect maxima function to count PLA signals across the entire image. The signal number was then divided by the number of nuclei to yield the signal per cell ratio for each image.

### Confocal and TIRF Microscopy Image Acquisition and Analysis

Immunofluorescently-stained neurons were imaged using a Leica SP8 inverted confocal microscope and Leica LAS X software, with sequential channel acquisition to minimize spectral overlap. Acquisition parameters such as laser power, detector gain, and pinhole size, were kept constant across experimental conditions within each experiment. Z-plane images were utilized for represented images of samples unless otherwise specified, then Z-plane images were selected at maximum intensity projections. Total internal reflection fluorescence (TIRF) microscopy was performed on fully differentiated living SH-SY5Y neurons (in #1.5 glass-bottomed plates in insulin-free Medium 3), using a Leica Thunder Live Cell 3D microscope and Hamamatsu W-VIEW GEMINI optical image splitter under controlled environmental conditions (37 °C, 5% CO₂), with identical acquisition settings applied across conditions. Image processing was limited to linear adjustments of brightness and contrast applied uniformly across entire images. Time-lapse images were generated from TIRF videos with ImageJ, and kymographs were generated with the KymoResliceWide ImageJ plugin. Analysis of protein colocalization using Pearson’s Correlation Coefficient from confocal images of IF staining was done via FIJI/ImageJ software using the Coloc2 plug-in. Region of interests (ROI) were applied when appropriate and Pearson’s Correlation Coefficient (R-Value) was determined by the plug-in to quantify colocalization between proteins of interests.

### Statistical Data Analysis

For quantitative comparisons of normally distributed measured values from two groups, we used Student *t*-tests with differences deemed significant if the calculated p-value <0.05. For multiple comparisons, we used one-way or two-way ANOVA followed by appropriate post hoc tests, with significance determined if p <0.05.

## DATA AND MATERIALS AVAILABILITY

Requests for further information and resources should be directed to and will be fulfilled by the lead contact, Dr. Ben Fogelgren. Proteomics data have been deposited at the MassIVE repository (MSV000100595) and is publicly available.

## ACKNOWLEDGMENTS

This work was supported by grants from the National Institutes of Health [R01DK117308, U54MD007601-38S1 to B.F.]; Hawaii Community Foundation [19ADVC-95450 to B.F.]; and the University of Hawaii at Manoa YIBR COBRE program [P20GM103457, Pilot award to B.F. and R.A.N.]. We are grateful to the University of Hawaii Cancer Center’s Microscopy, Imaging, and Flow Cytometry Core (NIH P30CA071789) for helping us with TIRF microscopy and live imaging of our fluorescent lines. We acknowledge the Proteomics Core Facility at the University of Arkansas for Medical Sciences and the IDeA National Resource for Quantitative Proteomics (NIH R24GM137786) for their services.

## AUTHOR CONTRIBUTIONS

C.B., G.P., R.S., M.O., R.A.N., and B.F. conceived and designed the research. C.B., G.P., R.S., H.K., and B.F wrote the manuscript. C.B., G.P., R.S., H.K., S.S., S.A., A.J.L., L.N., B.E.H., M.O., and B.F. executed experiments, including data collection and analysis. J.B.O., N.P., M.O., R.A.N, and B.F. interpreted results and provided key insights. All authors were involved in editing the manuscript and approving the final submitted version.

## DECLARATION OF GENERATIVE AI AND AI-ASSISTED TECHNOLOGIES

Portions of this manuscript were refined using ChatGPT to improve language clarity and readability. References and citation formatting were assisted by Liner AI. All experimental data, analyses, figures, and scientific interpretations were generated and verified by the authors, who take full responsibility for the accuracy and integrity of the work.

## SUPPORTING INFORMATION

### SI Methods

#### Proteomics

Protein samples were subjected to chloroform/methanol extraction and enzymatically digested with trypsin. Peptide samples were analyzed on an Orbitrap Exploris 480 mass spectrometer operated in data-independent acquisition (DIA) mode, using a 60-minute chromatographic gradient for each sample. Detailed LC–MS/MS parameters were determined by the facility according to their protocols. Raw mass spectrometry data were processed and searched using Spectronaut (Version 19; Biognosys) against the appropriate reference proteome database. Peptide and protein identifications were controlled at a false discovery rate (FDR) of 1%. Bioinformatic analysis was also performed at UAMS using MaxQuant and ProteiNorm. This workflow included quality control assessment, data normalization, and statistical analysis. Protein intensities were log2-transformed and normalized to correct for systematic variation across samples. Data quality was evaluated using standard quality control metrics. Differential expression analysis was performed within the analysis pipeline. The principal component (PC) plot and violin plot were generated using protein abundance data and statistical outputs provided by UAMS. The bar graphs and volcano plot were re-rendered in Graphpad Prism and functionally relevant proteins were consolidated together for comparative interpretation. These modifications were limited to visualization and did not affect raw data or quantitative analyses.

#### Antigen Retrieval

For the formalin-fixed paraffin embedded wildtype mouse brain sections (Amsbio, #MP-201-09-C57), antigen retrieval was done by deparaffinizing slides in 3 baths of ClearRite3 (Richard Allan Scientific, #6901) for 5 minutes each followed by rehydration of the slides in EtOH for 3 minutes in the following concentrations: 100%, 100%, 95%, 85%, 70%, and 50% with a final incubation in distilled water. Slides were then boiled in 1mM of EDTA buffer (Thermoscientific Chemicals, #J15694.AP) at 90°C for 20 minutes total. Slides remained in EDTA buffer and cooled for 35 minutes. Slides were then rinsed twice with 1X PBST for 5 minutes each, then incubated with 3% H_2_0_2_ (Fisher BioReagents, BP2633500) for 15 minutes at RT. Slides were rinsed with 1X PBST for 5 minutes each and dried followed by blocking at RT for 1 hour in 0.1% BSA (Sigma, #A7030-50G). From here, brain tissues were labeled with PLA conjugates based on manufacturer’s protocol.

## Supplemental Figures

**Figure S1.**
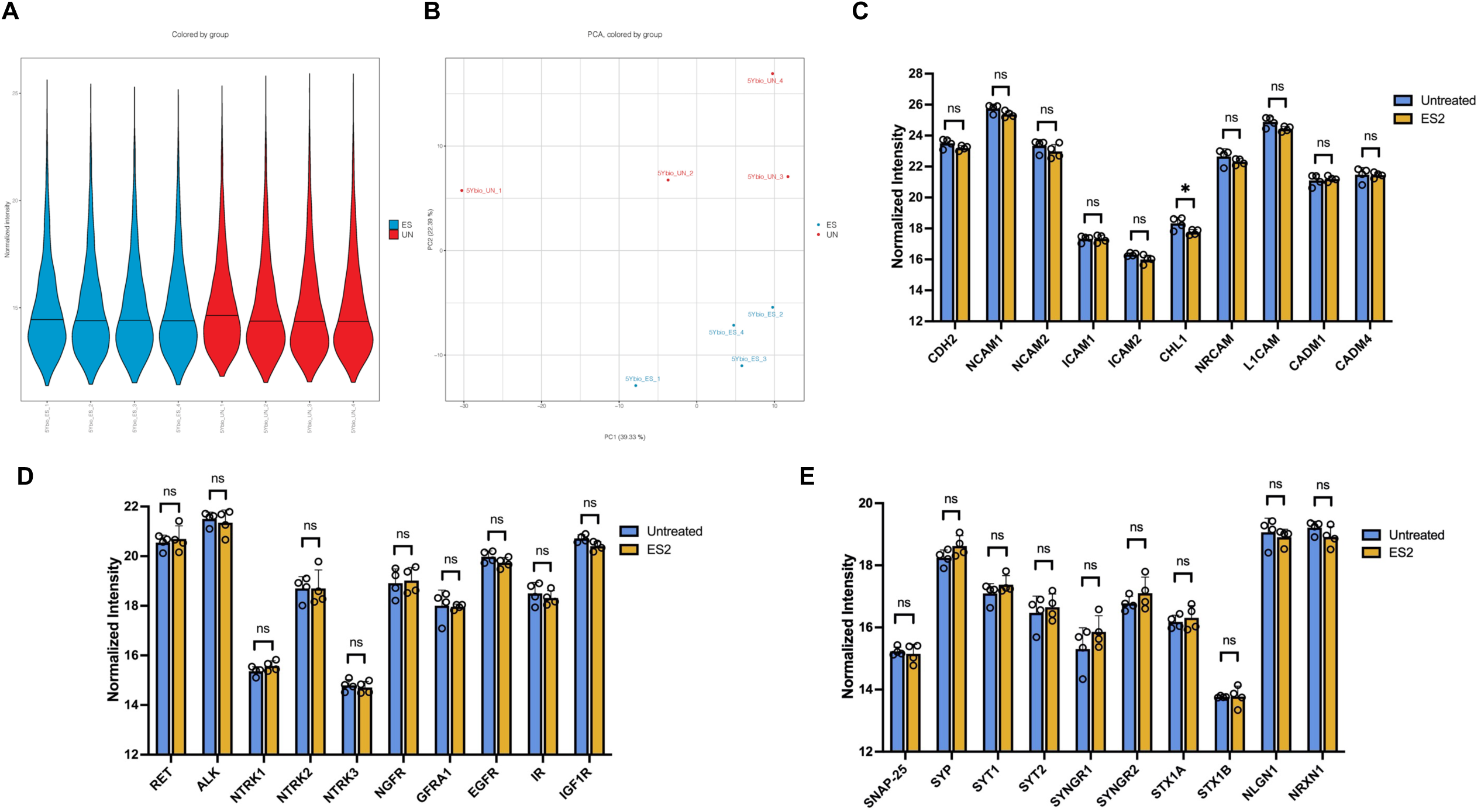
Proteomics screen quality control and representative groups of cell surface proteins unchanged in differential abundance after ES2 treatment. A. Violin plot showing consistent global normalization and distribution of log2-normalized protein intensities across each samples per treatment (red: untreated samples, blue: ES2-treated samples). B. Principal component analysis of log2-normalized protein intensities shows clear distinction between treated (blue) and untreated samples (red), with PC1 and PC2 explaining 61% of the total variance, indicating that the treatment is a major contributor to proteomic variation. C. Normalized intensity of biotinylated representative cell adhesion proteins, such as CDH2, NRCAM, CADM4, with most showing no significant changes after ES2 treatment. D. Normalized intensity of biotinylated representative plasma membrane signaling receptors, such as RET, NTRKs, IR, and other related neurotrophic proteins, each showing no significant changes after ES2 treatment. E. Normalized intensity of biotinylated representative neuronal synaptic proteins, such as SNAP-25, SYNGR1/2, NRXN1, each showing no significant changes after ES2 treatment. Comparison between two samples/conditions was performed using an unpaired Student’s t-test. Group values were compared with one-way ANOVA and corrected via post-hoc Tukey test. Significance value with ±SD shown as *(p<0.05), **(p<0.01), ***(p<0.001), **** (p<0.0001), and ns (not significant).

**Figure S2.**
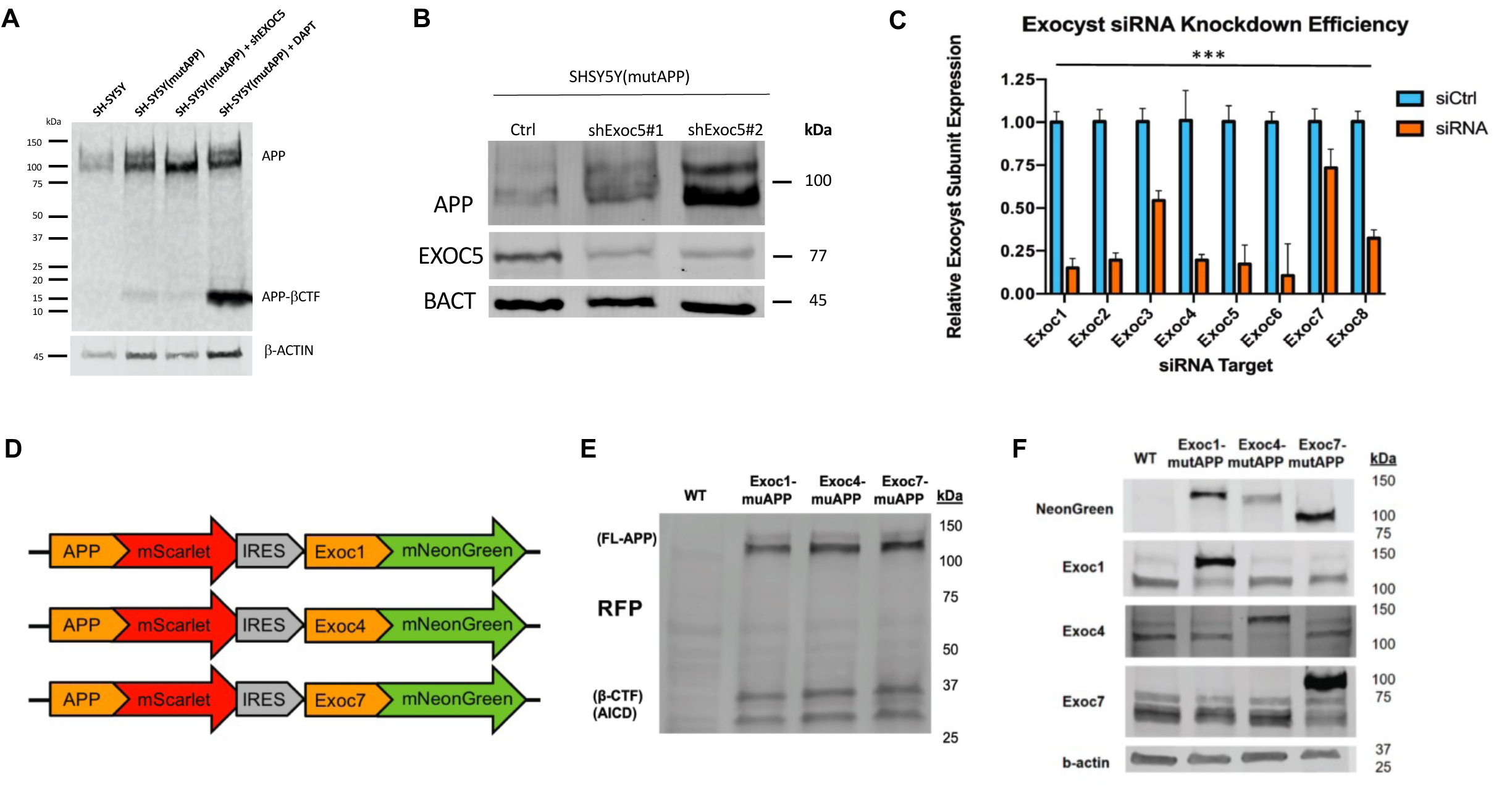
Validation of exocyst gene knockdown and fluorescently-labeled APP and EXOC5 transgenes in SH-SY5Y cells. A. Representative immunoblot image showing EXOC5 protein knockdown in two cell lines (shExoc5#1 and shExoc5#2). B. Quantitative real time PCR measurements demonstrated reduced exocyst gene expression in SH-SY5Y(mutAPP) cells after transfection with siRNAs against the individual eight exocyst genes. C. Schematic representation of three fluorescent transgene constructs co-expressing mutAPP fused with mScarlet on the C-terminus, and either EXOC1, EXOC4, or EXOC7 fused with mNeonGreen on the C-terminus. D. Validation of mutAPP-mScarlet expression in the three generated transgenic SH-SY5Y cell lines through immunoblotting with anti-RFP antibody. E. Validated expression of exocyst subunits (EXOC1,4,7) fused to mNeonGreen in the three generated transgenic SH-SY5Y cell lines through immunoblotting with anti-mNeonGreen and anti-exocyst antibodies.

**Figure S3.**
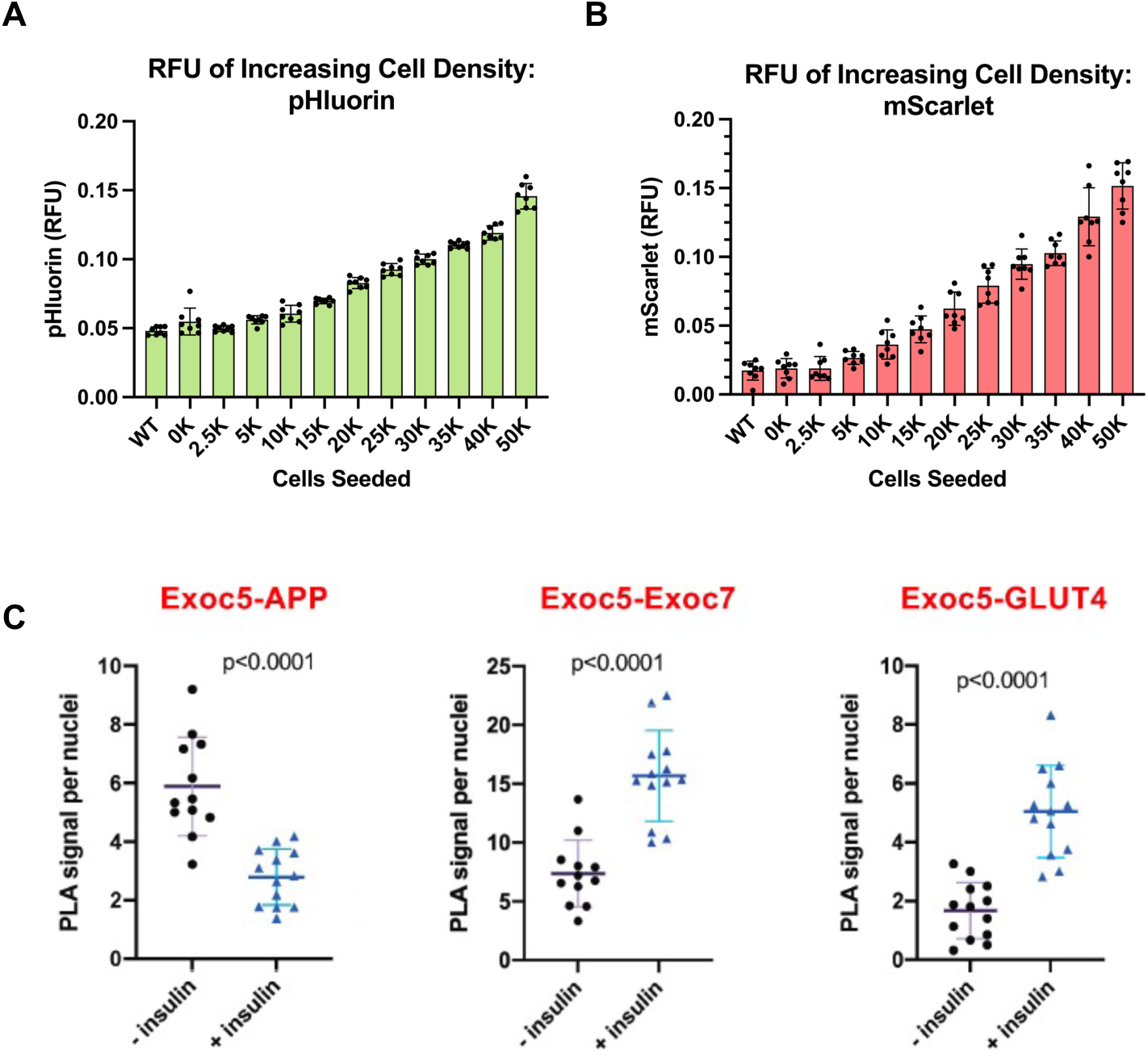
Initial microplate characterization of stable pHluorin-GLUT4-mScarlet expressing SH-SY5Y neurons, and PLAs of SH-SY5Y neurons in response to insulin. A. Representative images of EXOC5-APP PLA signals (red puncta) present throughout mouse pHCN dendrites and axon, as identified by immunostaining for MAP2 and TAU (both green, separate experiments), respectively, in insulin-starved condition (-Ins). B. Representative images of EXOC5-GLUT4 PLA signals (red puncta) observed in the soma and neurites of previously insulin-starved mouse pHCN after 15 minutes of 1 nM insulin addback. Scale bar= 10 µm C. Representative images of EXOC5-GLUT4 PLA signals (red puncta) showing increased associations in the soma and neurites of previously insulin-starved mouse pHCN after 15 minutes of 100 nM insulin addback. Scale bar= 5 µm A,B. Fluorescent microplate reading of differentiated SH-SY5Y neurons stably expressing the pHluorin-GLUT4-mScarlet fusion protein, confirming increasing both pHluorin and mScarlet signal with increasing cell seeding density. F. Fluorescent microplate reading of differentiated SH-SY5Y neurons stably expressing pHluorin-GLUT4-mScarlet showed pHluorin signal decreased when the pH of the cell medium was lowered, confirming pH sensitivity of pHluorin. G. Fluorescent microplate reading of differentiated SH-SY5Y neurons stably expressing pHluorin-GLUT4-mScarlet showed pHluorin/mScarlet RFU decreased with longer insulin starvation times. H. Comparison of before and after 15 minutes of 100nM insulin addback after varying lengths of insulin starvation. Longer starvation periods yielded greater significant increases after insulation stimulation. Statistical analysis was completed using a paired T-Test as the same biological replicates during starved conditions were measured after insulin treatment.

**Supplementary Table 1:** Primary and Secondary Antibodies used in this study.

| Target | Antibody/Clone (mono, poly) | Host | Application | Dilution | Vendor | Catalog # |
|---|---|---|---|---|---|---|
| β-actin | Monoclonal | Mouse | WB | 1:000 | ProteinTech | 5606 |
| ANKG | Polyclonal | Chicken | IF | 1:500 | Novus | NBP3-05549-50ul |
| APP | Monoclonal | Rabbit | IF, WB | 1:100, 1:1000 | Abcam | ab32136 |
| ATG5 | Monoclonal | Rabbit | WB | 1:1000 | Cell Signaling | D5F5U |
| BSN | Monoclonal | Mouse | IF | 1:100 | Abcam | ab82958 |
| EXOC1 | Polyclonal | Rabbit | WB | 17:40 | ProteinTech | 11690-1-AP |
| EXOC4 | Monoclonal | Mouse | WB | 1:1000 | Enzo | ADI-VAM-SV016 |
| EXOC5 | Monoclonal | Mouse | IF, WB | 1:100, 1:1000 | Santa Cruz | Sc-514802 |
| EXOC7 | Polyclonal | Rabbit | WB | 1:1000 | ProteinTech | 12014-1-AP |
| GLUT4 | Monoclonal | Mouse | IF, WB | 1:100, 1:500 | Santa Cruz | Sc-53566 |
| LAMP1 | Monoclonal | Rabbit | WB | 1:1000 | Cell Signaling | 9091 |
| LC3A/B | Monoclonal | Rabbit | WB | 1:1000 | Cell Signaling | 4445 |
| MAP2 | Polyclonal | Chicken | IF | 1:500 | Abcam | ab5392 |
| RAB11 | Monoclonal | Rabbit | WB | 1:1000 | Cell Signaling | 5589S |
| RAB5 | Monoclonal | Mouse | IF | 1:1000 | Cell Signaling | 46449S |
| RFP | Monoclonal | Mouse | WB | 1:1000 | Chromotek | 6g6 |
| Synaptophysin | Polyclonal | Goat | IF | 1:100 | R&D Systems | AF5555 |
| TAU | Polyclonal | Chicken | IF, WB | 1:500 | Abcam | ab75714 |
| **Selected Secondary Antibodies** |  |  |  |  |  |  |
| Antibody | Host: Target | Application |  | Dilution | Vendor | Catalog # |
| Alexa Fluor 488 (green)- Secondary Antibody | Goat: Chicken | IF |  | 1:1000 | ThermoFisher Scientific | A32931 |
| Alexa Fluor 594 (red)- Secondary Antibody | Goat: Chicken | IF |  | 1:1000 | ThermoFisher Scientific | A-11042 |
| Donkey Anti-Goat IgG H&L (Alexa Fluor® 647) preabsorbed | Donkey: Goat | IF |  | 1:1000 | Abcam | ab150135 |
| Licor Secondary (red) | Goat: Rabbit | WB |  | 1:10,000 | Licor | 926-68071 |
| IRDye® 680RD IgG Secondary Antibody | Donkey: Mouse | WB |  | 1:10,000 | Licor | 926-68072 |
| IRDye® 800CW IgG Secondary Antibody | Goat: Rabbit | WB |  | 1:10,000 | Licor | 926-32211 |

